# TIPs-VF: An augmented vector-based representation for variable-length DNA fragments with sequence, length, and positional awareness

**DOI:** 10.1101/2025.02.15.637782

**Authors:** Marvin I. De los Santos

## Abstract

The ability to accurately encode and represent genetic sequences in machine learning process is critical for advancements in biotechnology, specifically in genetic engineering and synthetic biology. Traditional sequence encoding method face significant limitations in handling sequence variability, maintaining reading frame integrity, and preserving biologically relevant features. This preliminary study presents TIPs-VF (Translator-Interpreter Pre-seeding for Variable-length Fragments), a simple and efficient encoding framework designed to address some of the key challenges in representing genetic sequences for machine learning. The results showed that TIPs- VF enables a variable-length sequence representation that retains biological context while ensuring the alignment of encodings with codon boundary, making it particularly suited for modular genetic construction. TIPs-VF demonstrated superior performance in truncation and fragmentation analysis, sequence homology detection, domain assessment, and splice junction identification. Unlike conventional methods that require fixed-length inputs, TIPs-VF dynamically adapts to sequence length variations, preserving essential features such as domain similarities and sequence motifs. Additionally, TIPs-VF improves open reading frame recognition and enhances the identification of vector parts and plasmid elements by unifying sequence embeddings with the three possible open reading frame. Overall, TIPs-VF offers a robust, biologically meaningful encoding framework that overcomes the constraints of traditional sequence representations by incorporating sequence, length, and positional awareness. The TIPs-VF encoding infrastructure is available at https://tips.logiacommunications.com.

## INTRODUCTION

The advancement of machine learning (ML) has revolutionized numerous fields, including biology and biotechnology. In recent years, ML algorithms have demonstrated remarkable capabilities in analyzing vast and complex datasets, providing insights that were previously unattainable through conventional methods. These advancements have enabled researchers to model biological systems, predict molecular interactions, and optimize biotechnological processes with unprecedented accuracy and efficiency. In particular, the intersection of machine learning and genomics has opened new frontiers in understanding and manipulating genetic information (1).

The applications of artificial intelligence (AI) in the field of genetics and genomics have paved the way for groundbreaking developments in genetic engineering and synthetic biology. AI algorithms have been employed to predict gene functions (2), identify genetic variations linked to diseases (3), and design synthetic genes with specific functionalities (4). These innovations are driving the development of personalized medicine, sustainable agriculture, and bio-based manufacturing. For such applications, AI relies heavily on the ability to effectively process and analyze genetic sequence data, highlighting the importance of accurately representing this information in computational models (5).

The representation of genetic sequences is a critical factor in applying machine learning to genomics. DNA sequences, composed of four nucleotide bases—adenine (A), thymine (T), cytosine (C), and guanine (G)—must be translated into formats that ML models can interpret. Traditional methods, such as one-hot encoding and k-mer frequency analysis, have been used to numerically represent DNA sequences (6). However, the high dimensionality and redundancy of these representations often pose challenges for machine learning applications, particularly when dealing with large genomic datasets (7).

Despite the progress made, there is a growing need for more sophisticated numerical representations of genetic sequences. Current approaches vary widely, ranging from simple binary encodings (8) to advanced embeddings derived from natural language processing techniques (9). While these methods have facilitated significant breakthroughs, they often struggle to capture the full complexity and hierarchical nature of genomic data. Furthermore, many existing representations are limited by their computational inefficiency or their inability to preserve important biological features, such as sequence motifs and structural properties.

A major shortcoming of current encoding schemes is their inability to effectively represent key sequence features, sequence length, base position, and reading frames in a unified manner. One-hot encoding, for example, results in highly sparse and high-dimensional representations that are computationally expensive. K-mer-based approaches, while compact, often fail to preserve sequence order and positional information, unless an overlapping approach is applied, which significantly increases the resource requirement. Additionally, embeddings inspired by natural language processing techniques may struggle with the inherent non-overlapping nature of biological sequences, leading to information loss and reduced interpretability. These limitations pose significant challenges in terms of efficiency, computational resources, and overall effectiveness in extracting meaningful biological insights from genomic data (10).

Addressing these gaps in encoding schemes is essential for advancing genetic engineering and synthetic biology. By developing novel representations that balance compactness, fidelity, and computational efficiency, researchers can unlock new possibilities in genome design, synthetic pathway optimization, and precision medicine. In this paper, the TIPs-VF (Translator-Interpreter Pre-seeding for Variable-length Fragments) encoding technique is introduced. The augmentation of genetic sequence representation of variable-length DNA fragments was explored. Additionally, the ability of TIPs-VF to analyze and predict DNA fragmentation, sequence homology, sequence motif, and splice junctions was investigated.

## METHODOLOGY

### Data acquisition

#### TIPs-VF validation

The cDNA sequences or coding DNA sequences (CDS) of genes in chromosome 21 were sourced from the ENSEMBL BioMart database (https://www.ensembl.org/biomart/martview/). The sequences were manually or programmatically examined to ensure data integrity. Any data containing characters other than A, T, C, and G were removed during validation to rule out the effects of sequence misrepresentation. The data did not undergo uniform-length preprocessing.

#### Fragmentation analysis

Specific genomic regions of chromosomes 1, 2, and 3 (124M, 121M, and 99M, respectively) were obtained from the NCBI GenBank (https://www.ncbi.nlm.nih.gov/genbank/). These genomic regions were trimmed to uniform 50,000-nucleotide lengths and further truncated at the 5’ and/or 3’ ends, generating subsequences of 12,500-nucleotide long. This fragmentation resulted in truncated sequences with the same sizes but different sequence compositions.

#### Sequence homology analysis

The complete genome sequences of SARS-CoV, SARS-CoV-2, MERS-CoV, influenzaviruses (Alphainfluenza and Betainfluenza), HIV, HPV, and measles viruses were obtained from the NCBI Virus database (https://www.ncbi.nlm.nih.gov/labs/virus/vssi/#/). Sequences not in the conventional A, T, C, G format were replaced with ‘N.’

#### Sequence motif analysis

Since the TIPs-VF encoding was designed for synthetic biology and cloning purposes, sequence motif analysis was performed using various cloning and expression vectors. Plasmid maps and sequences for CRISPR experiments, mammalian expression, yeast manipulation, and plant recombinant experiments were obtained from SnapGene (www.snapgene.com). Any sequences not in the conventional A, T, C, G format were replaced with ‘N.’

#### Splice junction analysis

The DNA fragments from chromosomes 1 and 2 (as described in Fragmentation analysis) were used for splice junction analysis. The sequences were randomly fragmented into 1,500-nucleotide segments with 1,000 iterations, resulting in non-identical sequence fragments. Each fragment from chromosome 1 was randomly spliced with a fragment from chromosome 2, ensuring no fusion fragments contained redundant sequences. Four experimental setups were conducted: (1) a fusion pair containing a gap junction, (2) an inverse fusion pair containing a gap junction, (3) a fusion pair without a gap junction, and (4) an inverse fusion pair without a gap junction. The gap junction sequence was identical for all experiments (Table 1).

**Table 1.**
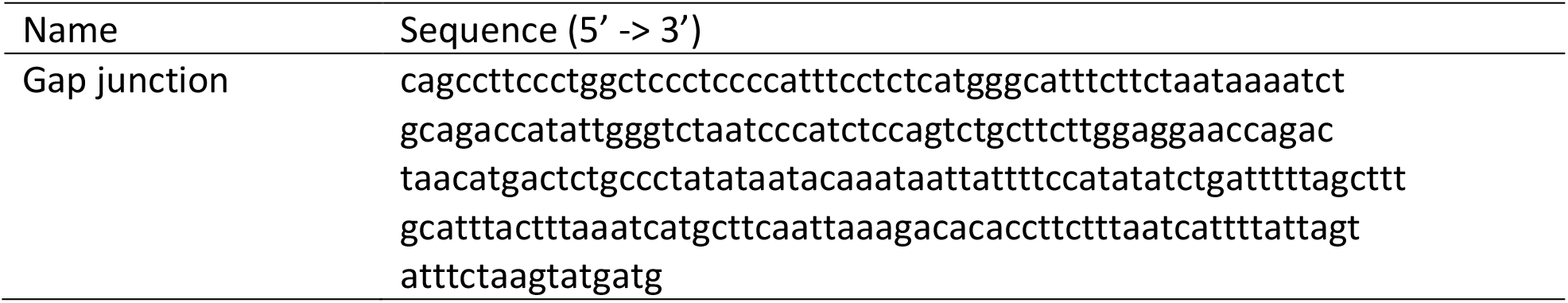
Gap junction sequence used in the splice junction analysis.

### Data preprocessing

Upon downloading, the sequences were inspected for FASTA formatting. Any sequences that failed to meet the required standards (e.g., missing labels, >50% of the sequence being ‘N,’ or less than 100 bp in length) were excluded. Non-A, T, C, G sequences were either replaced with ‘N’ or corrected by referencing an NCBI assembly. A Python-based preprocessing pipeline was implemented to remove duplicate entries and filter out sequences that did not meet the above standards.

### Representation of genetic sequences

The genetic sequences were numerically represented using three encoding methods. The first utilized DNABERT (5), a model specifically adapted for biological sequence data. The second encoding framework utilized a 6-mer representation, where two approaches were applied: 1) non-overlapping 6-mers, where sequences were divided into independent 6-mer units, and 2) a sliding window 6-mer method, which generated overlapping 6-mers to capture positional relationships.

The third representation method involved the development of TIPs-VF encoding (Figure 1). TIPs- VF was initially designed for generative augmentation of DNA sequences in genetic engineering and synthetic biology applications (e.g., construction of multi-specific antibodies). It aims to provide a representation logic that supports critical workflows in DNA fragment manipulation and feature extraction, including sequence characteristics (e.g., start codons), position, and length.

**Figure 1.**
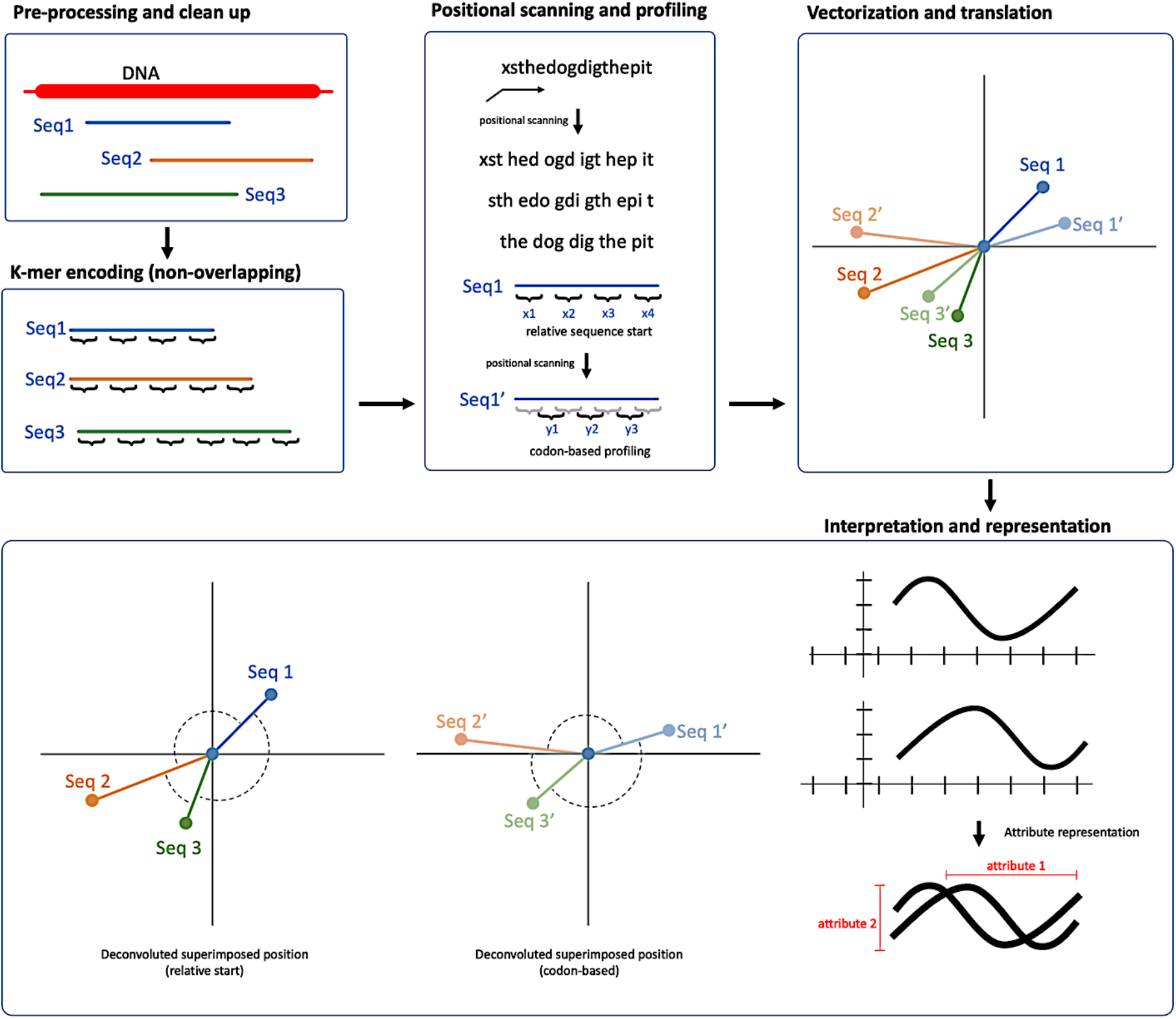
Overview of the design and representation of genetic sequences using the TIPs-VF (Translator-Interpreter Pre-seeding for Variable-length Fragments) encoding scheme. DNA sequences were pre-processed and cleaned to extract target sequences, followed by non-overlapping k-mer (initially by 6-mer) encoding, positional scanning, and codon-based profiling. The 6-mer units were translated and vectorized by calculating the similarity between each unit in the vector space. The sequences were then interpreted and represented based on the closeness and directional alignment of their attributes.

Current DNA sequence encoding techniques often fail to fully capture DNA attributes such as the relative position of fragments, a feature that is typically obscured in multidimensional representations. TIPs-VF is based on a non-overlapping k-mer (6-mer) representation that is independent of frequency association. This means that it does not depend on the frequency of specific k-mer units, which often lack positional information. Additionally, TIPs-VF operates in a non-overlapping parameter, making it computationally efficient and requiring lower resources (e.g., storage) for representation.

### Neural network implementation

The TIPs-VF encoding scheme was further evaluated through implementation in a neural network framework using Keras (11). The neural network architecture included an input layer for the encoded representations, multiple fully connected hidden layers with ReLU activation functions, and a softmax output layer for classification tasks. The model was compiled using Mean Squared Error (MSE) as the loss function and trained on scaled input data. All analyses were conducted in the Google Colab environment using Python, with key libraries including TensorFlow, scikit-learn, and matplotlib ensuring efficient workflow execution.

### Visualization of embeddings

After implementation in a deep neural network through Keras, the embeddings of the encoded representations were visualized using three dimensionality reduction techniques: PCA, t-SNE, and UMAP. Principal Component Analysis (PCA) was employed to reduce dimensionality while retaining variance. The t-Distributed Stochastic Neighbor Embedding (t-SNE) method projected high-dimensional embeddings into two-dimensional space to highlight local structure and relationships. Lastly, the Uniform Manifold Approximation and Projection (UMAP) method was applied to uncover global and local patterns, providing complementary insights to PCA and t-SNE. The visualization methods used in each figure are indicated in the figure captions and/or labels.

### Heatmap analysis

Pairwise distances among the embeddings generated from PCA, t-SNE, and UMAP were computed using the Euclidean distance metric. A heatmap representation was generated to visualize these distances and identify patterns in embedding similarities. Hierarchical clustering was applied to the distance matrices using Ward’s method, and dendrograms were constructed to illustrate the hierarchical relationships among the embeddings. All visualizations were implemented using Seaborn and Matplotlib in Python.

### TIPs-VF encoding availabilty

The TIPs-VF encoding scheme is available at https://tips.logiacommunications.com via a backend JavaScript implementation. A Python version is available on Google Colab with private access, upon reasonable request. The datasets and implementation of TIPs-VF are available on Github (https://github.com/mahvin92/TIPs-VF).

## RESULTS

### Representation of variable-length sequences using TIPs-VF

To test the hypothesis that TIPs-VF can be used to represent variable-length DNA fragments with a sequence-aware feature, we compared the TIPs-VF-based vector embeddings of chromosome 21 genes (in varying lengths) with k-mer frequency encoding using overlapping and non-overlapping representations. As a baseline, DNABERT was used to plot the sequences in a vector space to assess the absence of random clustering (to rule out false positives in the analysis), which is expected since DNABERT is dependent on contextual similarities (5) and frequencies (12), as shown in Figure 2a. Our initial results showed that k-mer frequency representation lacked sequence awareness (data not shown), likely due to the absence of a significant consensus sequence in chromosome 21 genes. To rule out the influence of random sequence misrepresentation, some genes were fragmented while retaining sequence similarities from the original gene sequence (e.g., SIK1 was fragmented into seven subsequences of varying lengths). Both non-overlapping and overlapping k-mer frequency representations failed to cluster fragments with similar or consensus sequences in different dimension reduction analyses (PCA, t-SNE, and UMAP), as shown in Figures 2b and 2c.

**Figure 2.**
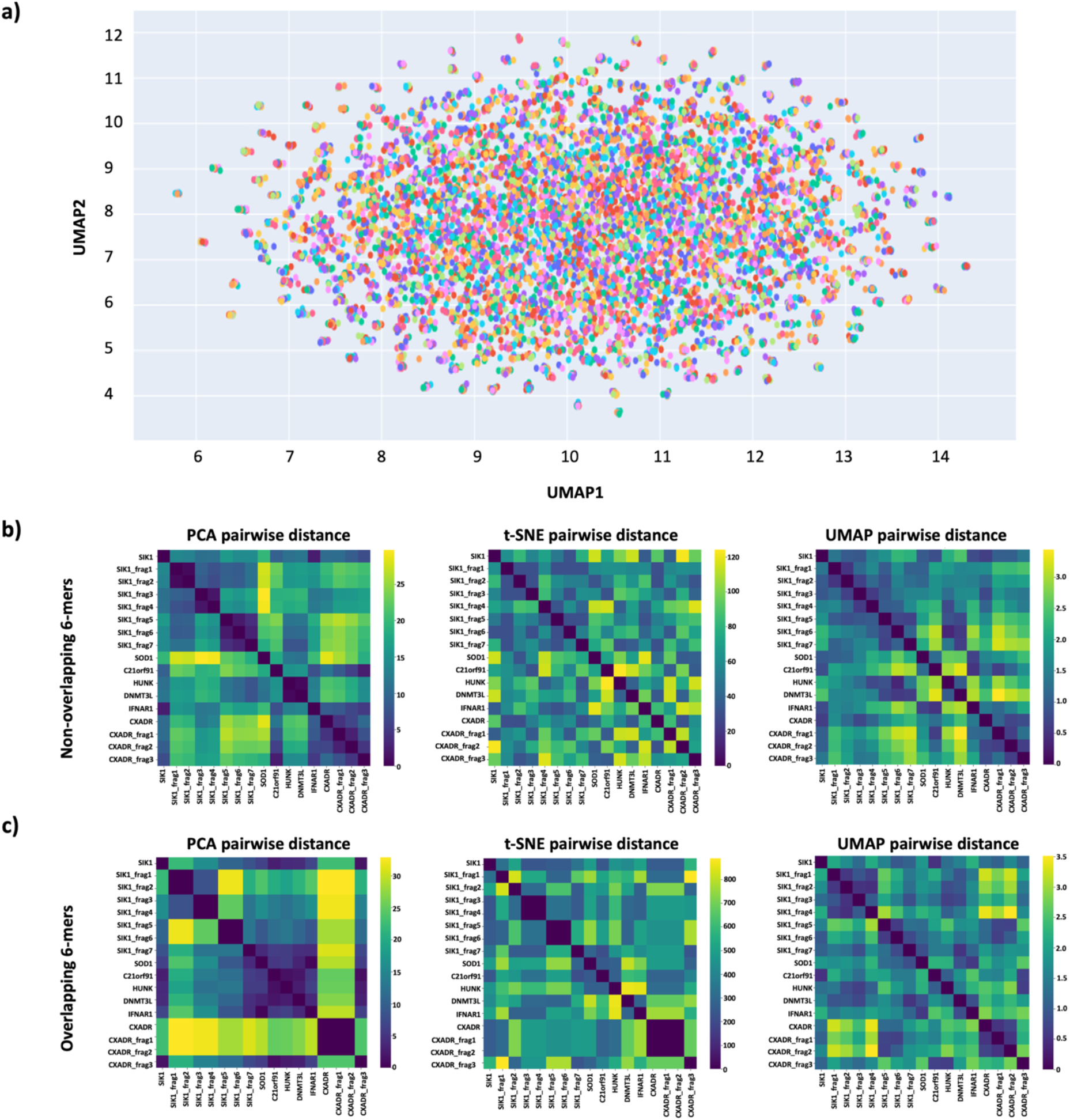
Representation of gene sequences in chromosome 21 using DNABERT and k-mer frequency through non-overlapping and overlapping encodings. **a**) Visualization of chromosome 21 gene embeddings generated by DNABERT using UMAP. **b-c**) Heat map analyses of the embedding pairwise distances for chromosome 21 genes represented by non-overlapping (**b**) and overlapping (**c**) 6-mer encodings following neural network implementation. Pairwise distances for fragmented and unfragmented sequences (only selected genes/fragments are shown) were calculated for each dimensionality reduction analysis (PCA, t- SNE, and UMAP, respectively).

Pairwise gene alignment scoring revealed the expected gene consensus or similarity patterns of the analyzed and fragmented sequences. The SIK1 gene and its seven fragments exhibited strong pairwise gene alignment scores (Figure 3a). The same input sequences were represented using TIPs-VF, which initially considered three scanning positions (that encode the three start positions of a possible reading frame) of the bases in each k-mer unit, as shown in Figure 3b. After representation, the TIPs-VF attributes were scored for similarity, and the results showed that for each attribute (attributes 1 and 2), the score of each representation with respect to their base position correlated strongly for each gene (e.g., scans 1–3 of the SIK1 gene in attribute 1 had scores of 0.99, 0.89, and 0.94, respectively) and for each related fragment (e.g., SIK1 had scan attribute scores in the range of 0.89–0.99, and SIK1 fragments 2–4 had scores within the same range), as shown in Figure 3c. This indicates that the representation of gene sequences and fragments by TIPs-VF efficiently captures sequence data, such as base position and codon information.

**Figure 3.**
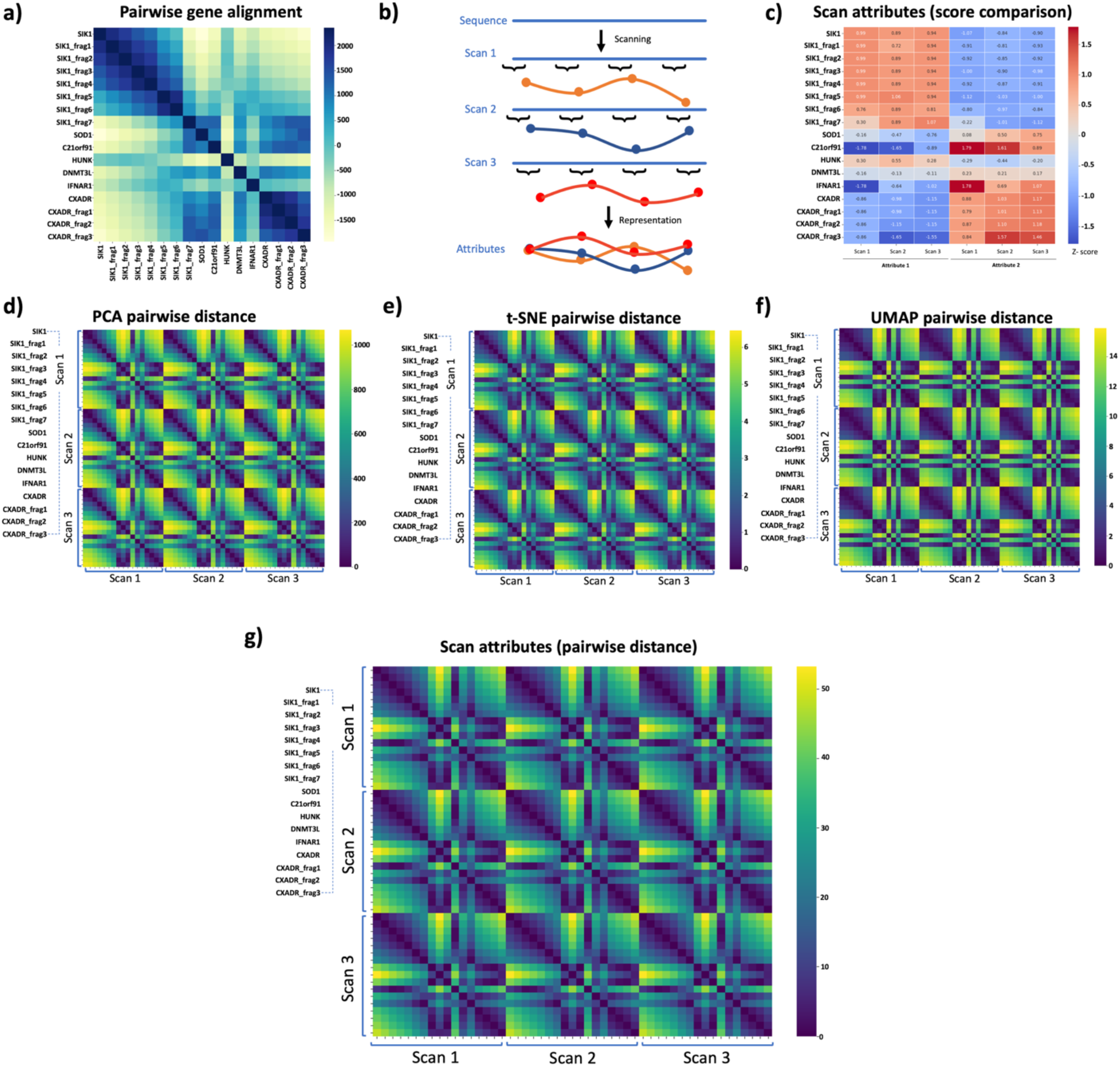
Representation of variable-length DNA fragments (chromosome 21 genes) using TIPs-VF. **a**) Heat map analysis of the pairwise gene alignment scores for fragmented and unfragmented chromosome 21 sequences, indicating the degree of similarity. **b**) Schematic of the three scan positions corresponding to the relative start positions of an open reading frame. The TIPs-VF encoding scheme is designed to simultaneously represent all three scanning attributes. **c**) Comparison of the scan attributes between fragmented and unfragmented sequences, revealing that TIPs-VF captures sequence similarity patterns in gene alignment scoring. **d–f**) Heat map analyses of the embedding pairwise distances for fragmented and unfragmented sequences (only selected genes/fragments are shown), calculated using PCA, t-SNE, and UMAP, respectively. **g**) Heat map analysis of the pairwise distances for the scores of the three scans in each analyzed sequence.

The implementation of TIPs-VF application in genetic sequence representation resulted in vector embeddings that captured the sequence similarities of gene fragments. Pairwise distance analysis of the vector embeddings of gene fragments showed that TIPs-VF was able to represent the fragments with high sequence similarities (e.g., strong pairwise distances between the SIK1 gene and its seven fragments). Indeed, for all dimension reduction analyses, the pairwise distances of all fragments captured the same pattern observed in pairwise gene alignment scoring (Figures 3d–3f). This result initially conveys the sequence-aware feature of TIPs-VF encoding and suggests its potential use for BLAST-free alignment of gene sequences with variable lengths. Lastly, to assess whether the TIPs-VF attributes were altered during neural network implementation, the pairwise distance pattern of each sequence representation was obtained. The results showed that TIPs-VF representation was implemented successfully without significant alteration in the encoded sequence data (Figure 3g).

### Uniform representation of base codon by TIPs-VF

To further test the position awareness of TIPs-VF in representing genetic sequences, the comparative patterns of different scans corresponding to the three start positions of a possible reading frame were analyzed. Figure 3c shows that the encoded attributes of each scan were comparable. However, the results also showed that TIPs-VF relies on similarity coverage for sequence identification. For example, the SIK1_frag7 fragment, which had the lowest similarity score to the SIK1 gene (Figure 3a), also had the lowest attribute score comparison in some of the scans (Figure 3c). Nonetheless, one advantage of TIPs-VF encoding compared to pairwise gene alignment scoring is that the sums of the three scan attributes for each fragment derivative from a gene sequence correlated positively (range: 2.65–2.99), which could be more reliable than the weighted comparison used in gene alignment scoring. The reliability of TIPs-VF over gene alignment scoring needs to be further assessed in larger dataset testing.

### Truncation and fragmentation analysis

Given the indications that TIPs-VF can augment the representation of sequence similarities and scan (e.g., reading frame) or base position of DNA fragments, the hypothesis that TIPs-VF can be used for BLAST-free alignment using longer sequences was tested. Three genomic regions spanning 20K–30K nucleotides fragments in chromosomes 1, 2, and 3 were truncated to 15,000- nt, followed by fragmentation into uniform lengths of 12,500-nt but with varying levels of internal sequence similarity. TIPs-VF representation resulted in chromosome-specific clustering of the fragments (Figure 4a). Pairwise embedding distance analysis revealed three distinct heatmap patterns corresponding to the genomic fragments in chromosomes 1–3 (Figure 4b). Sequence similarity assessment found that fragments in chromosome 1 had a higher degree of similarity with fragments in chromosome 3 than in chromosome 2. This finding is corroborated by the closer distances of fragment clustering in UMAP embedding visualization and shorter pairwise distances in the heatmap analysis. These observations align with dendrogram clustering, as shown in Figure 4c. These results signify the applicability of TIPs-VF for BLAST-free alignment and potentially for sequence-based gene identification.

**Figure 4.**
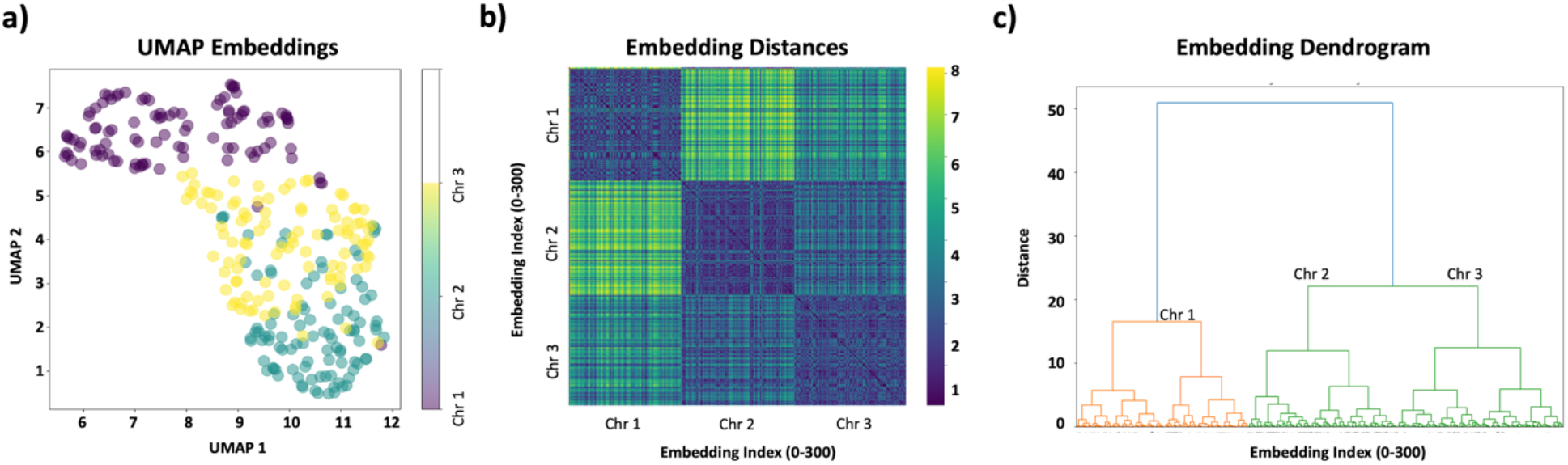
Representation of chromosomal fragments and truncated sequences by TIPs-VF. **a**) Visualization of fragment embeddings from chromosomes 1, 2, and 3 using TIPs-VF, analyzed with UMAP. **b**) Heat map analysis of the embedding pairwise distances for fragments from the three chromosomal regions. **c**) Dendrogram clustering of the calculated embedding pairwise distances.

### Sequence homology analysis

With the assumption that TIPs-VF can be used for BLAST-free alignment, we next determined its application for sequence homology identification. We conducted this experiment by representing the complete genome sequences of Coronaviruses (SARS-associated coronavirus, SARS-CoV-2, and MERS-related coronavirus), Influenzaviruses (Alphainfluenzavirus and Betainfluenzavirus), and Non-respiratory viruses (HIV, HPV, and Measles viruses). TIPs-VF representation resulted in sequence homology clustering, as expected, where Coronaviruses, Influenzaviruses, and Non-respiratory viruses formed distinct clusters (Figure 5a). Pairwise embedding distance analysis revealed three distinct heatmap patterns corresponding to the three viral groups (Figure 5b).

**Figure 5.**
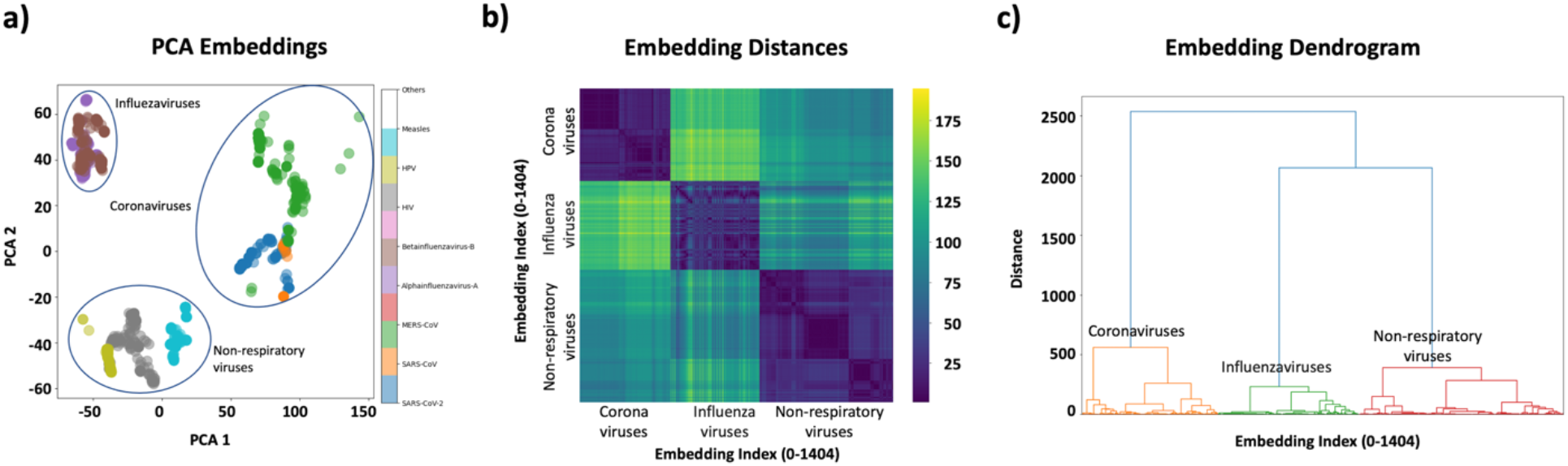
Representation of different viral genome sequences by TIPs-VF. **a**) Visualization of genome sequence embeddings from various viruses using TIPs-VF, analyzed with PCA. **b**) Heat map analysis of the embedding pairwise distances for the viral genome sequences. **c**) Dendrogram clustering of the calculated embedding pairwise distances.

Sequence homology assessment found that Coronaviruses formed non-overlapping and distinct clusters from Influenzaviruses and Non-respiratory viruses. Further in-group sequence homology assessment revealed that different taxonomic classifications of the viruses were captured. For example, the distance embedding within Coronaviruses revealed that SARS-CoV-2 sequences had a distinct pairwise distance score from other Coronavirus species, which was also evident in the sub-clustering of SARS-CoV-2 data points. These observations align with dendrogram clustering, as shown in Figure 5c. These results suggest that the TIPs-VF encoding scheme has potential applications in sequence homology comparisons, such as gene prediction for recombinant analysis or taxonomic classification.

### Sequence motif assessment

With evidence that TIPs-VF can predict and weigh sequence similarities or diversities, its ability to identify and differentiate cloning or expression-related motifs in plasmids and other recombinant vectors for downstream genetic engineering and synthetic biology applications was examined. DNA sequences of cloning and expression vectors for mammalian, plant, yeast, and CRISPR workflows available on SnapGene were represented using TIPs-VF. Neural network implementation resulted in closed-loop clusters that nearly resembled ‘domain fragments’ due to overlapping data points (Figure 6a). Further analysis revealed close associations between representations of recombinant vectors for mammalian and CRISPR applications, which is expected given the widespread applications of CRISPR vectors for mammalian genome editing (Figure 6b). Dendrogram analysis revealed four distinct clusters, likely corresponding to the four vector classifications/types (Figure 6c).

**Figure 6.**
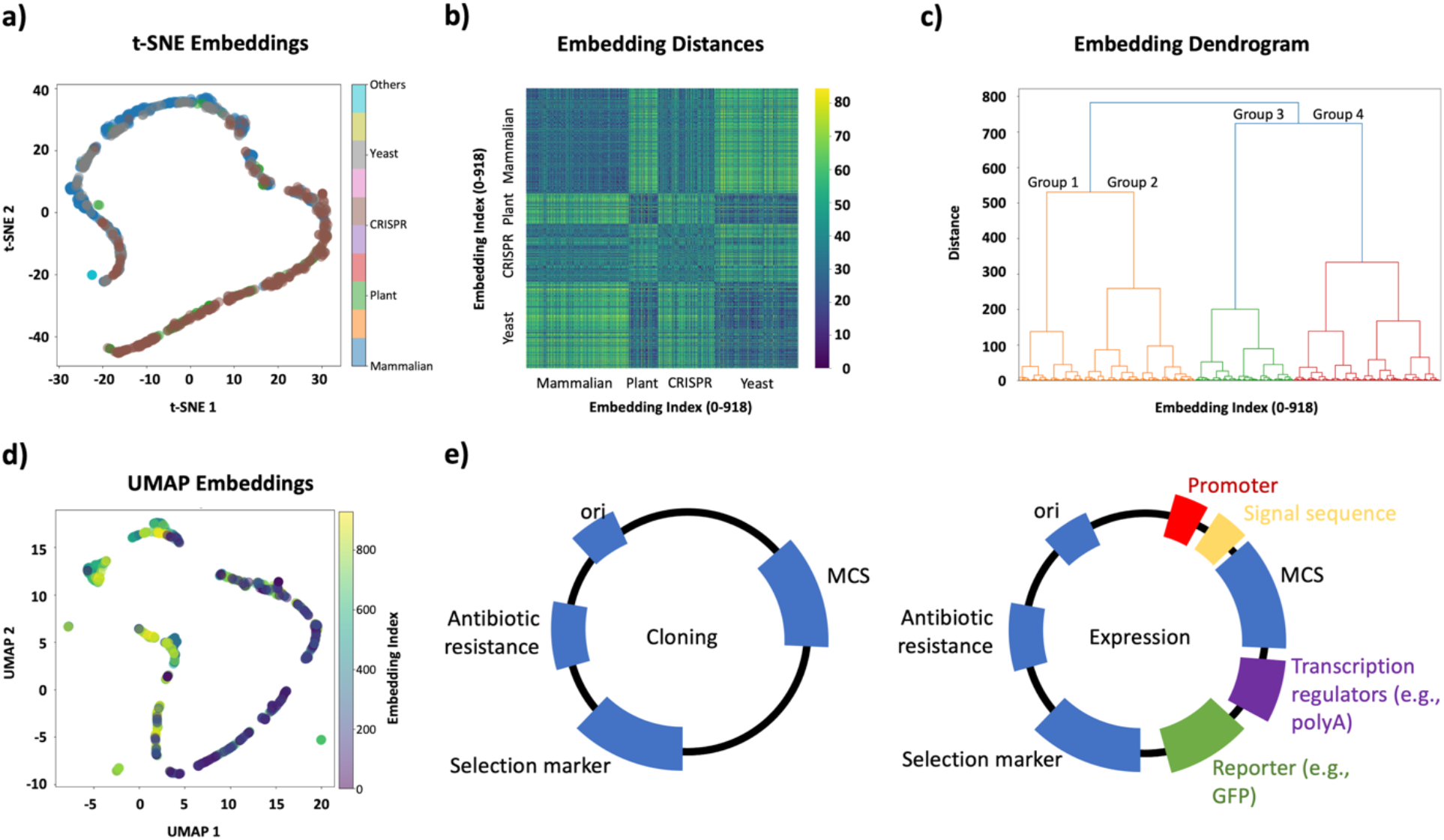
Representation of different recombinant vector sequences by TIPs-VF. **a**) Visualization of vector sequence embeddings from various cloning and expression plasmids using TIPs-VF, analyzed with t-SNE. **b**) Heat map analysis of the embedding pairwise distances for the recombinant vectors. **c**) Dendrogram clustering of the calculated embedding pairwise distances. **d**) Visualization of vector sequence embeddings from various cloning and expression plasmids using TIPs-VF, analyzed with UMAP. **e**) Graphical representation of the canonical and common components of cloning and expression vectors.

To further investigate the proposed ‘domain fragments,’ a different dimensionality reduction analysis was performed. UMAP visualization of the TIPs-VF embedding revealed more distinct fragmented clusters, which are likely indicative of the motif or domain features of various plasmid vectors. In this study, TIPs-VF representations were limited to entire vector sequences rather than individual vector components. However, as shown in Figure 6d, certain sequence attributes of mammalian vectors are shared with other vector types. This is expected, as vector lineages can often be remodeled or re-engineered to fit specific applications (e.g., swapping yeast-specific promoters for mammalian promoters).

Although it is premature to conclude that the observed domain fragments directly correspond to sequence homologies resulting from vector sequence similarities, this study hypothesizes that TIPs-VF, driven by sequence and base position similarities, can capture vector domains and regulatory regions in vectors that share high sequence consensus, including sequence motifs (Figure 6e). Further evaluation through motif-specific experiments is required to validate this hypothesis.

### Splice junction analysis

Finally, the ability of TIPs-VF to augment sequence junction recognition in neural networks was examined. Curated fusion genes from published literature were first analyzed; however, clustering was unsuccessful due to the lack of consensus sequences or low sequence similarity at the splice junctions (data not shown). To account for this limitation, a new analysis was performed using synthetically generated fusion genes with splice gaps of uniform length and without gaps (Figure 7a).

**Figure 7.**
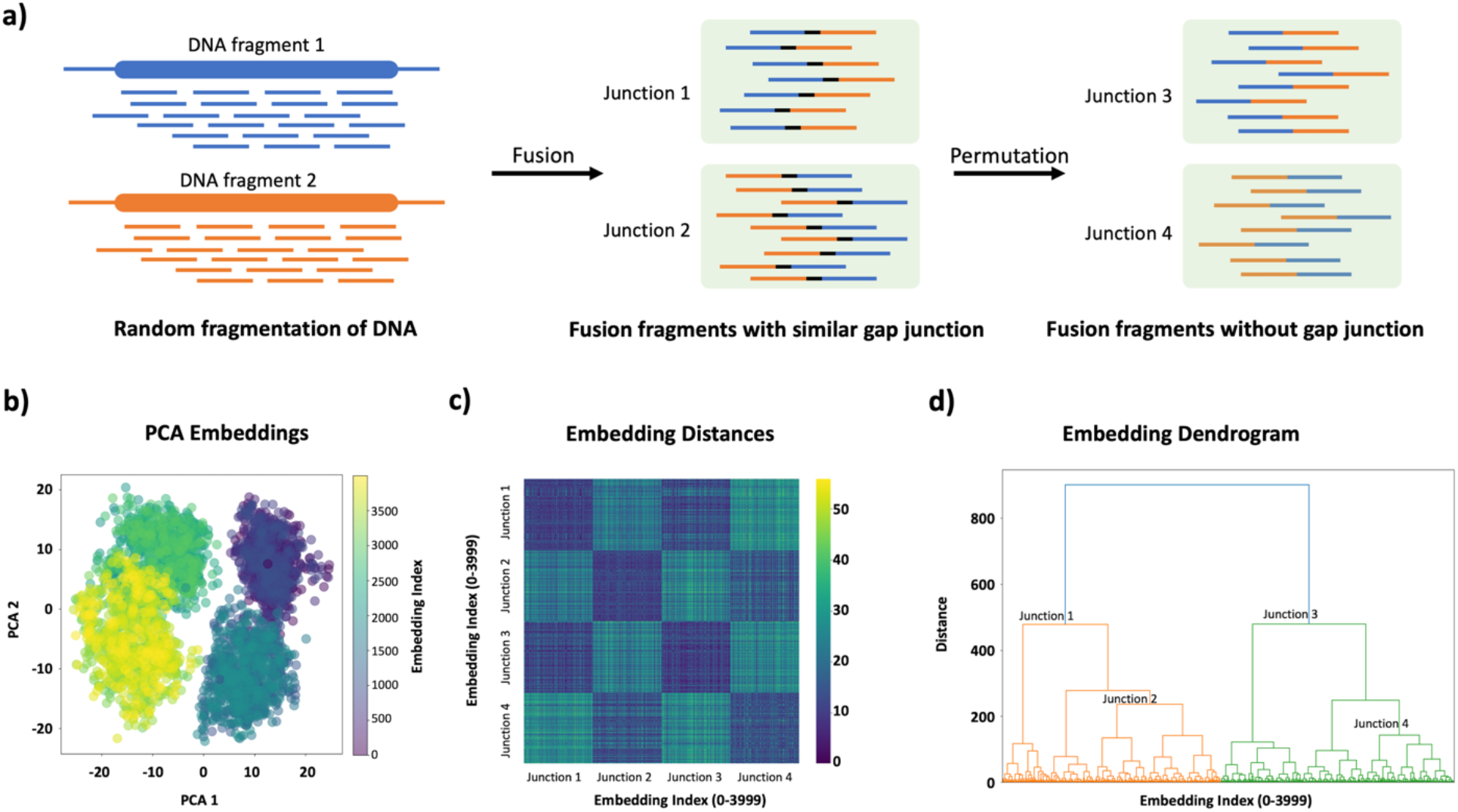
Representation of different fusion/spliced fragments by TIPs-VF. **a**) Overview of the splice junction analysis. Two DNA fragments from chromosomes 1 and 2 were randomly fragmented and fused in different pairing orders with a gap junction. A permutation without a gap junction was included to represent a group lacking a consensus gap sequence. **b**) Visualization of fusion fragment embeddings from different junction groups using TIPs-VF, analyzed with PCA. **c**) Heat map analysis of the embedding pairwise distances for the different junction groups. **d**) Dendrogram clustering of the calculated embedding pairwise distances.

TIPs-VF representations of the synthetically generated fusion genes resolved into distinct clusters corresponding to different junction groups (Figure 7b). Interestingly, despite having similar k-mer frequency distributions, junctions 1 and 2 clustered separately due to the insertion of a 250-nt gap sequence, which caused a shift in the reading frame. This suggests that TIPs-VF may enhance the representation of reading-frame-aware genetic features. In contrast, junctions 3 and 4, which lacked gaps, exhibited overlapping features—an expected result since the fusion of the two DNA fragments did not alter the reading frame.

Pairwise embedding distance analysis revealed four distinct heatmap patterns corresponding to the four junction groups (Figure 7c). Additionally, junction 1 showed a higher degree of similarity to junction 3, which is expected given that their only difference was the presence or absence of the 250-nt gap sequence. A similar pattern was observed for junctions 2 and 4. These findings were further corroborated by dendrogram clustering (Figure 7d).

## DISCUSSION

The ability to augment, manipulate, modify, and enhance biological features has always been the holy grail of biotechnology. Today, two interconnected fields have emerged as powerful tools to achieve this goal: genetic engineering and synthetic biology. Genetic engineering involves the precise manipulation of genetic material to enhance or alter the genotypic or phenotypic traits of living organisms. These fields are computationally intensive and biologically resource-heavy disciplines (13-16). They integrate advancements in computational tools, data analysis, and predictive modeling to design and validate biological constructs.

Despite their transformative potential, genetic engineering and synthetic biology face significant challenges, particularly in biocomputational application and design complexity. These challenges have led to the integration of artificial intelligence (AI) and machine learning (ML) to enhance research and development (17-18). Today, AI tools can now predict gene-editing outcomes, optimize metabolic pathways, and simulate the behavior of synthetic biological systems. For example, reinforcement learning algorithms have been used to optimize the output of a co-culture bioprocess (19), while generative adversarial networks (GANs) have been found to aid in generating optimized sequences for creating highly diverse functional proteins (20).

Despite these advances, traditional sequence encoding schemes still present major limitations in genetic engineering and synthetic biology. Many commonly used methods, such as one-hot encoding, k-mer frequency analysis, and positional encoding, struggle with the inherent variability of biological sequences (6,10). These conventional encoding strategies often require fixed-length inputs, if not combined with other approaches, making them less suitable for representing sequences of varying lengths, such as those found in plasmid vectors, fusion genes, and regulatory elements to name a few. This constraint limits the ability to accurately capture essential sequence features, such as fragment identity, structural motifs, and evolutionary homologies. Moreover, most conventional encoding schemes fail to incorporate reading frame- aware representations (21), leading to a loss of biologically meaningful information when analyzing sequences that undergo translational shifts or alternative splicing events.

TIPs-VF overcomes these limitations by enabling a variable-length sequence representation that retains the biological context of genetic elements. Unlike fixed-dimensional embeddings, TIPs-VF dynamically adapts to sequence length variations, ensuring that truncations and fragmentations are accurately represented without loss of essential information (22). This advantage is particularly relevant for analyzing plasmid domains, where functional regions may be partially conserved across different vector types. By preserving fragment-specific sequence integrity, TIPs- VF enhances the resolution of domain truncation and fragmentation analysis, making it a valuable tool for studying modular genetic components.

Another critical advantage of TIPs-VF is its ability to uniformly represent base codons in a way that maintains a unified reading frame for vectorization. Traditional methods often treat sequences as mere character strings, neglecting the structural importance of codon boundaries (23). This oversight can obscure crucial genetic features when analyzing translated sequences. TIPs-VF mitigates this issue by aligning sequence embeddings to the reading frame, allowing for improved recognition of open reading frames (ORFs), regulatory elements, and translational control regions. This capability is especially beneficial for studies focusing on gene expression regulation, protein coding potential, and functional annotation of synthetic constructs.

The results of this study demonstrate the practical advantages of TIPs-VF across multiple sequence analysis tasks. TIPs-VF effectively distinguishes domain fragments and truncated sequences, preserving sequence context while maintaining the relationships between functionally relevant elements. This capability is crucial for designing synthetic vectors and modular genetic elements with predictable functionality. Next, unlike traditional methods that rely on global sequence alignments, TIPs-VF captures local sequence similarities, even among distantly related genetic elements. This feature allows for more accurate detection of homologous regions and evolutionary relationships, which is essential in synthetic biology applications involving recombination and domain shuffling.

Furthermore, TIPs-VF enhances motif recognition by incorporating sequence position information, which is often lost in conventional frequency-based methods. This improved representation enables more precise identification of regulatory motifs, transcription factor binding sites, and conserved sequence patterns in engineered constructs. Lastly, this preliminary study also revealed that TIPs-VF improves the resolution of splice junction clustering by preserving sequence context and reading frame continuity. This advantage is particularly relevant in transcriptomic analyses and fusion gene detection, where accurate junction mapping is critical for understanding alternative splicing and gene fusion events.

In the proposed context of pre-seeding the representation of genetic sequences, TIPs-VF serves as modular encoding that can serve both as a foundational representation or conduit for subsequent computational tasks. This modularity allows TIPs-VF to be seamlessly integrated into various machine learning workflows, enhancing its adaptability across different computational frameworks and research applications.

While TIPs-VF offers significant advantages, there are several limitations to this study. Firstly, the dataset used was relatively small, which may limit the generalizability of the findings across diverse biological contexts. Future work should explore larger and more diverse datasets to validate the robustness of TIPs-VF in different genomic applications. Additionally, while TIPs-VF provides a strong framework for sequence representation, deeper integration with machine learning models, particularly deep learning approaches, was not explored in this study. Further research is needed to assess how TIPs-VF-encoded sequences perform in predictive tasks, such as classification and functional annotation using neural networks. Another limitation is that AI training and validation applications were not conducted, which could provide deeper insights into the scalability and adaptability of TIPs-VF in computational genomics. Furthermore, this study did not account for potential biases in sequence representation, which could influence downstream analyses. Future iterations should examine bias mitigation strategies to ensure fair and accurate representation of genetic elements across different biological datasets. Lastly, while TIPs-VF improves upon traditional encoding schemes, its computational efficiency and scalability in handling extremely large genomic datasets remain to be fully assessed.

## CONCLUSION

In conclusion, the findings suggest that TIPs-VF provides a robust and biologically relevant approach to encoding genetic sequences. By addressing the shortcomings of traditional encoding schemes, TIPs-VF facilitates more accurate sequence representation, enhances computational analysis, and supports the design and interpretation of genetic constructs in synthetic biology and genetic engineering. However, further validation and integration with deep learning applications are necessary to fully realize its potential in AI-driven biological research.

